# Pre-Columbian treponemes clarify worldwide spread of treponematosis

**DOI:** 10.1101/2024.01.15.575648

**Authors:** Gueyrard Mattéo, Pontarotti Pierre, Drancourt Michel, Abi-Rached Laurent

## Abstract

Syphilis dramatically hit Europe at the end of the fifteen century before spreading to other continents. Yet the origin of the sudden pandemic in the Old World remains debated, in particular because the leading Columbus hypothesis of a New World origin of historical syphilis in Europe lacks paleomicrobiological confirmation. Here we screened a worldwide set of >1,700 ancient humans and identified ancient *Treponema pallidum* strains in two pre-Columbian child sacrifices from Tlatelolco, Mexico. Over 12,000 *Treponema*-specific reads were recovered to define a novel *Treponema pallidum* ancient population: *Treponema pallidum* str. *tlatelolcoensis*. Phylogenetics show that this population displays ancestral features but also bears the genetic building blocks of disease-causing modern *Treponema pallidum* subspecies, hence demonstrating how pre-Colombian Americas were the source of worldwide spread of treponematosis.

## Introduction

Syphilis started ravaging Europe at the end of the fifteen century^1, 2^ and is still a worldwide disease over five hundred years later, with modern pandemic strains that trace back to a common ancestor in the mid-twentieth century^3^ having caused an estimated 6.3 million cases in 2016^4^. Although the first complete genome of the syphilis spirochete was characterized 25 years ago^5^, and dozens of additional *Treponema* genomes followed, the antiquity, sources, and dynamics of diffusion of human treponematoses in modern populations in Europe remains uncertain. Part of the challenge is that the *T. pallidum* subspecies that cause venereal syphilis (*T. pallidum* subsp. *pallidum)* and non-venereal yaws (*T. pallidum* subsp. *pertenue*) and bejel (*T. pallidum* subsp. *endemicum*) remain undistinguishable by morphology and antigenicity^6, 7^, and can only be identified through study of multiple genetic markers and near whole-genome sequencing^7, 8, 9^. In addition, only a limited number of investigations convincingly yielded ancient *Treponema pallidum* (*T. pallidum*) complex in Europe and Mexico, and none of those could be identified as unambiguously pre-Columbian^10, 11, 12^. Hence the leading Columbus hypothesis of a New World origin of historical syphilis in Europe^13, 14^ lacks paleomicrobiological confirmation.

Here we searched for *Treponema pallidum* strains in a worldwide set of >1,700 ancient human genomes and identified two such cases from 14^th^-15^th^ century Mexico^15^. These ancient genomes prove that *Treponema pallidum* genomes existed in pre-Columbian Americas and provide an unprecedented opportunity to define what role Christopher Columbus’ travels played in worldwide spread of syphilis.

## Results

### Two ancient Treponema genomes from pre-Columbian Tlatelolco, Mexico

To search for ancient *Treponema pallidum* genomes, we screened a worldwide set of over 1,700 ancient human genomes (Supplementary Figs. 1-2) with five *Treponema*-specific probes (see Methods). Two positive cases were identified in the remains of children who died in Tlatelolco city, Mexico in AD 1325-1520, at ages 4-6 (individuals IF #9 and IF #11 in ref^15^). Genomewide investigation isolated over 12,000 unique reads through a stringent 3-step procedure designed to capture all specific *Treponema* reads: 8,922 reads for IF #9 and 3,458 reads for IF #11 (Supplementary Fig. 3). The reads displayed the typical features of ancient DNA (Supplementary Fig. 4) and were then mapped against a reference *T. pallidum* genome to cover 38% (428kb; IF #9) and 17% (193kb; IF #11) of that reference (Fig. 1b,d) with a homogenous distribution (Fig. 1a, c).

**Fig. 1.**
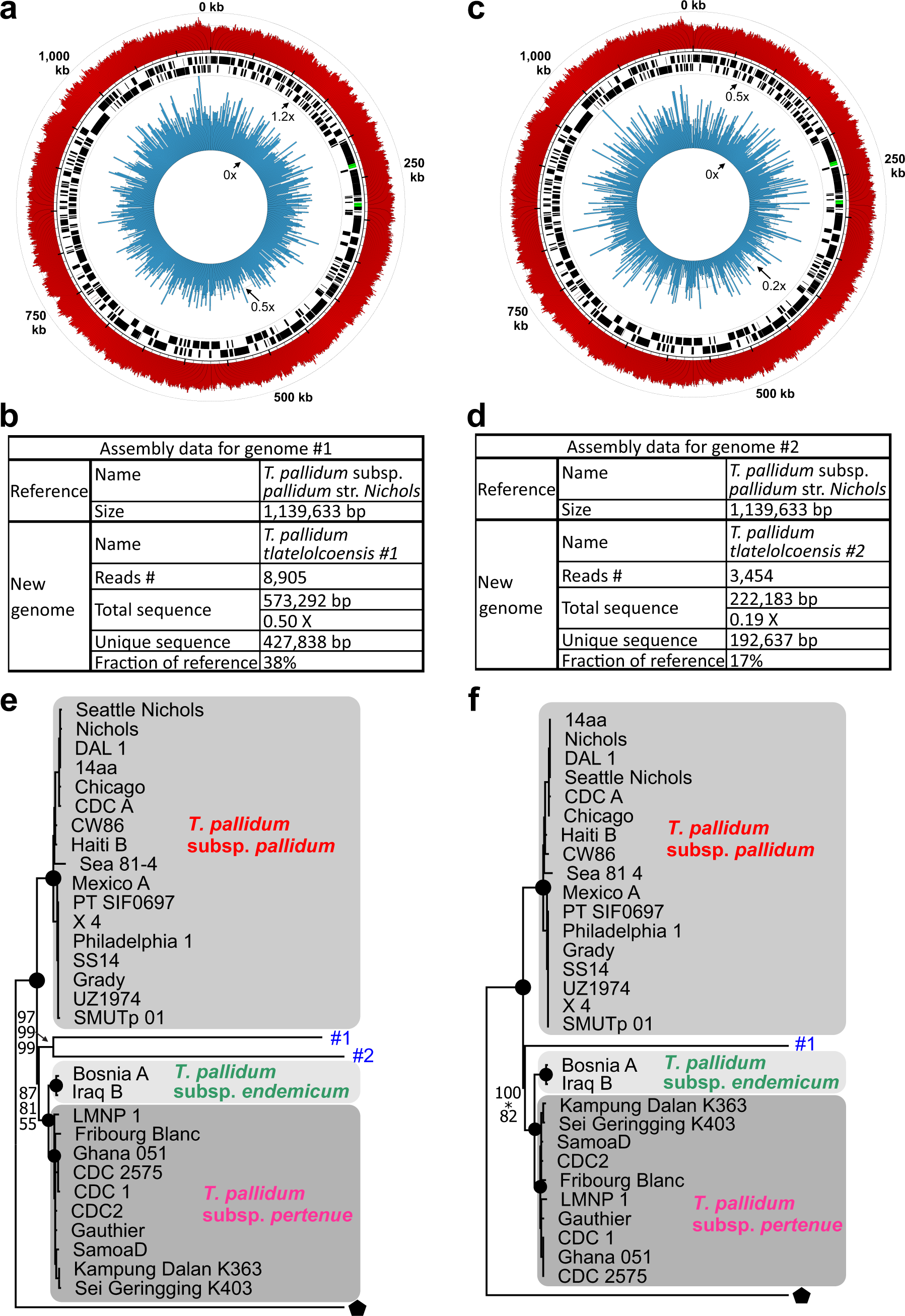
*Treponema pallidum* str. *tlatelolcoensis* is a basal, pre-Columbian *T. pallidum* strain. (**a** to **d**) Characteristics of the *T. pallidum* str. *tlatelolcoensis* genomes from individuals IF #9 (**a** and **b**) and IF #11 (**c** and **d**). (**a** and **c**) Circleators plot for the novel genomes with read coverage (internal layer; minimum, average and maximum coverage are given) plotted against GC content (outer layer) and gene content (two middle layers) from the reference (*T. pallidum* subsp. *pallidum* strain *Nichols*). (**b** and **d**) Statistics for the genome assemblies. (**e** and **f**) *T. pallidum* str. *tlatelolcoensis* is a basal *T. pallidum* strain. Full-genome phylogenetic analysis on the common segments of 32 (**e**) or 31 (**f**) *Treponema* genomes using neighbor-joining (NJ), parsimony, and maximum-likelihood (ML) methods. The NJ tree topology was used for the display, with a midpoint rooting. Bootstrap support is given for six (**f**) or seven (**e**) nodes (from top to bottom: ML, parsimony, NJ). Circles at nodes indicate bootstrap support of 100 with all methods. *, bootstrap support <50. Black pentagons, *T. paraluiscuniculi* outgroup.

### The Tlatelolco genomes are new members of the T. pallidum complex

To investigate the relationships between the two Tlatelolco genomes and other *Treponema* genomes, we first performed a genomewide phylogenetic analysis with the raw sequences using three methods (Fig. 1e). All three methods indicate that the two strains represent a monophyletic group that is a sister group to that formed by *T. pallidum* subsp. *pertenue* and *T. pallidum* subsp. *endemicum*. Yet, the two Tlatelolco genomes appear relatively distinct on the phylogenetic tree and display a raw divergence of ∼1.4% (Supplementary Fig. 5). Taking into account the differences that are due to the cytosine deamination of the ancient DNA (Supplementary Fig. 4) cuts this divergence to an estimated maximum of 0.86% (Supplementary Fig. 5). This is still a relatively high level and so it is possible these genomes are indeed distinct despite originating from the same ancient city. Consistent with this possibility, the two genomes display the same differences at 21 positions than those observed between the modern *Trepanoma* genome sequences used in the alignment.

Hence the Tlatelolco genomes represent the first two Pre-Columbian sequences of the *T. pallidum* complex. Because they also represent a new clade, we named them *T. pallidum* str. *tlatelolcoensis* #1 (IF #9) and #2 (IF #11).

### T. pallidum str. tlatelolcoensis carries distinct phylogenetic signals

To further analyze the *T. pallidum* str. *tlatelolcoensis* sequences, we then focused on just the largest sequence of the two so that the region in common with the other sequences would shift from ∼72kb (Fig. 1e) to ∼403kb (Fig. 1f). The first noticeable difference is that while the position of the sequence in the phylogenetic tree does not change, phylogenetic support does and increases with two methods (NJ, ML) but decreases with one (parsimony). While this could be just phylogenetic noise, this could also be indicative of underlying divergent signals in the dataset (see comment in methods) and to test this possibility, we did a simple split of the dataset in two halves and analyzed them independently (Supplementary Fig. 6). This analysis shows that for the first half of the dataset (Supplementary Fig. 6a and 6b), the methods diverge significantly, as assessed by a Shimodaira-Hasgawa test of alternative phylogenetic hypotheses that does reject the NJ and parsimony trees (Supplementary Fig. 6d; alpha=0.05).

Because visual inspection of the alignment hinted at the presence of divergent signals in different regions, we set to test the homogeneity of the phylogenetic signal by dividing the whole genome into segments. For this analysis, we used windows of 15kb after checking that this would lead to segments with an average of at least 10 differences between the *T. pallidum* subspecies (Supplementary Fig. 7), which should ensure good resolution of the relationships between the groups. Interestingly, this analysis reveals distinct patterns regarding the position of *T. pallidum* str. *tlatelolcoensis* comparing to the three modern *T. pallidum* strains: for 7 of the 27 segments (∼26%), its split is anterior to that of the three subspecies, for another 11 segments (∼41%), its split is anterior to that of two the three subspecies, and for four segments (∼15%) the split is associated with just one subspecies (Supplementary Fig. 8 and Fig. 2). To confirm this phylogenetic signal, we concatenated the segments with the same individual signal for the three largest groups (Fig. 2). This analysis shows strong and consistent phylogenetic signal between the methods for the position of the *T. pallidum* str. *tlatelolcoensis.* To further test this signal, we also conducted Shimodaira-Hasegawa tests of alternative phylogenetic hypotheses (Supplementary Fig. 9) and for two of the three patterns, these tests significantly reject the alternative tree topologies: for the clustering with the two subspecies *T. pallidum* subsp. *pertenue* and *T. pallidum* subsp. *endemicum* (alpha=0.001; Supplementary Fig. 9c) and for the clustering with *T. pallidum* subsp. *pallidum* (alpha=0.05; Supplementary Fig. 9b).

**Fig. 2.**
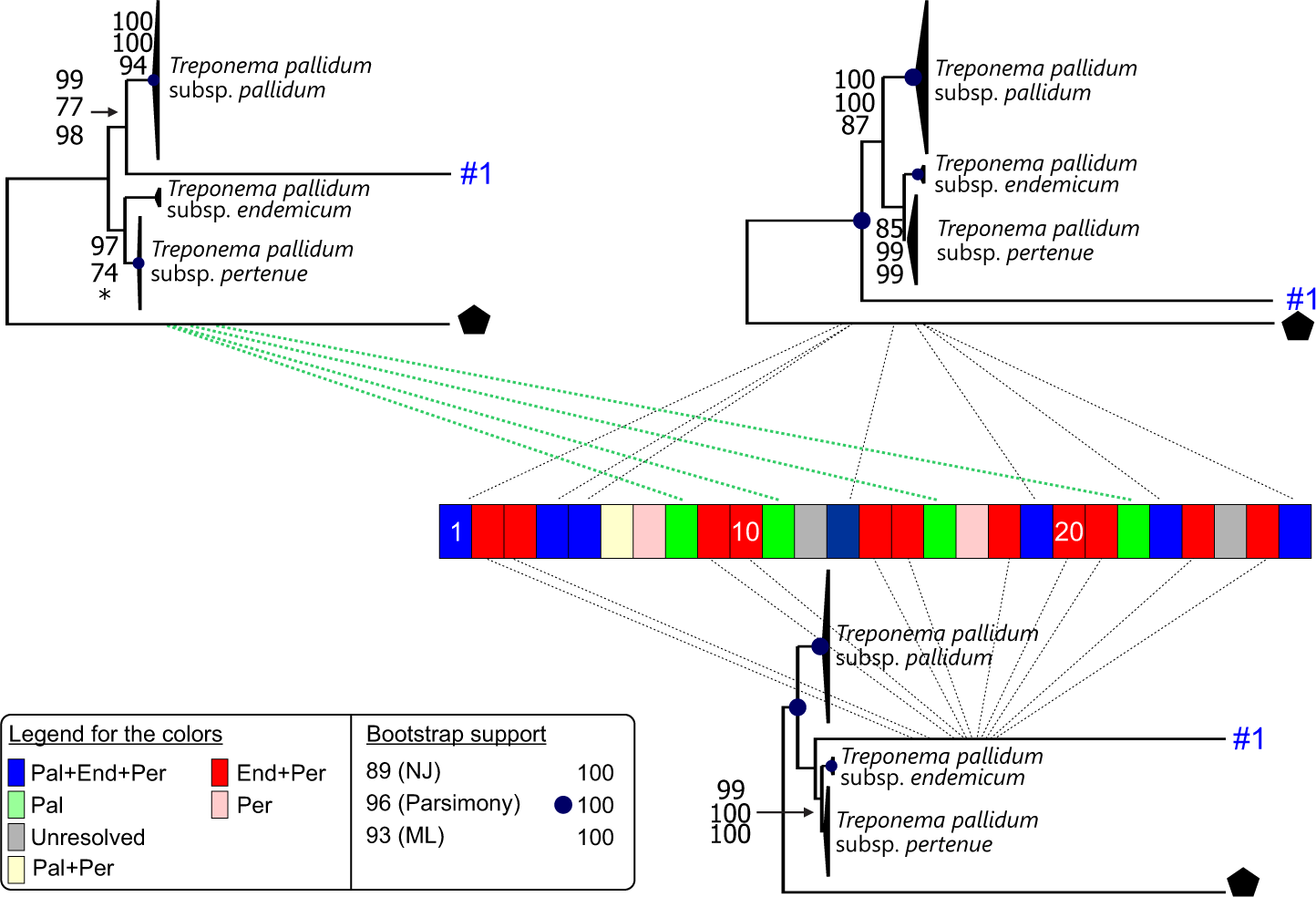
*T. pallidum* str. *tlatelolcoensis* displays distinct phylogenetic patterns in its relationships to modern *T. pallidum* subspecies. The genome sequence of *T. pallidum* str. *tlatelolcoensis* was divided in 27 segments of 15kb each and phylogenetically compared to the corresponding sequences of 30 other *Treponema* genomes using NJ, parsimony, and ML methods (Supplementary Fig. 8). The central bloc summarizes orthology to *T. pallidum* str. *tlatelolcoensis* for each segment using the colour scheme displayed in the bottom left corner. Pal, *T. pallidum* subsp. *pallidum*. End, *T. pallidum* subsp. *endemicum*. Per, *T. pallidum* subsp. *pertenue*. ‘Concatenated’ phylogenetic analyses were conducted for the three largest groups of segments: results are displayed above and below the central bloc and, together with statistical tests of topologies (Supplementary Fig. 9) confirm that these segments have distinct phylogenetic signals.

This analysis hence shows that *T. pallidum* str. *tlatelolcoensis* represents a genome with distinct phylogenetic signals. This pattern is consistent with *T. pallidum* str. *tlatelolcoensis* having evolved in pre-Columbian America as part of the ancestral *T. pallidum* population that experienced the transition from a single common ancestor to the three ancestors of the three *T. pallidum* subspecies (Fig. 3).

**Fig. 3.**
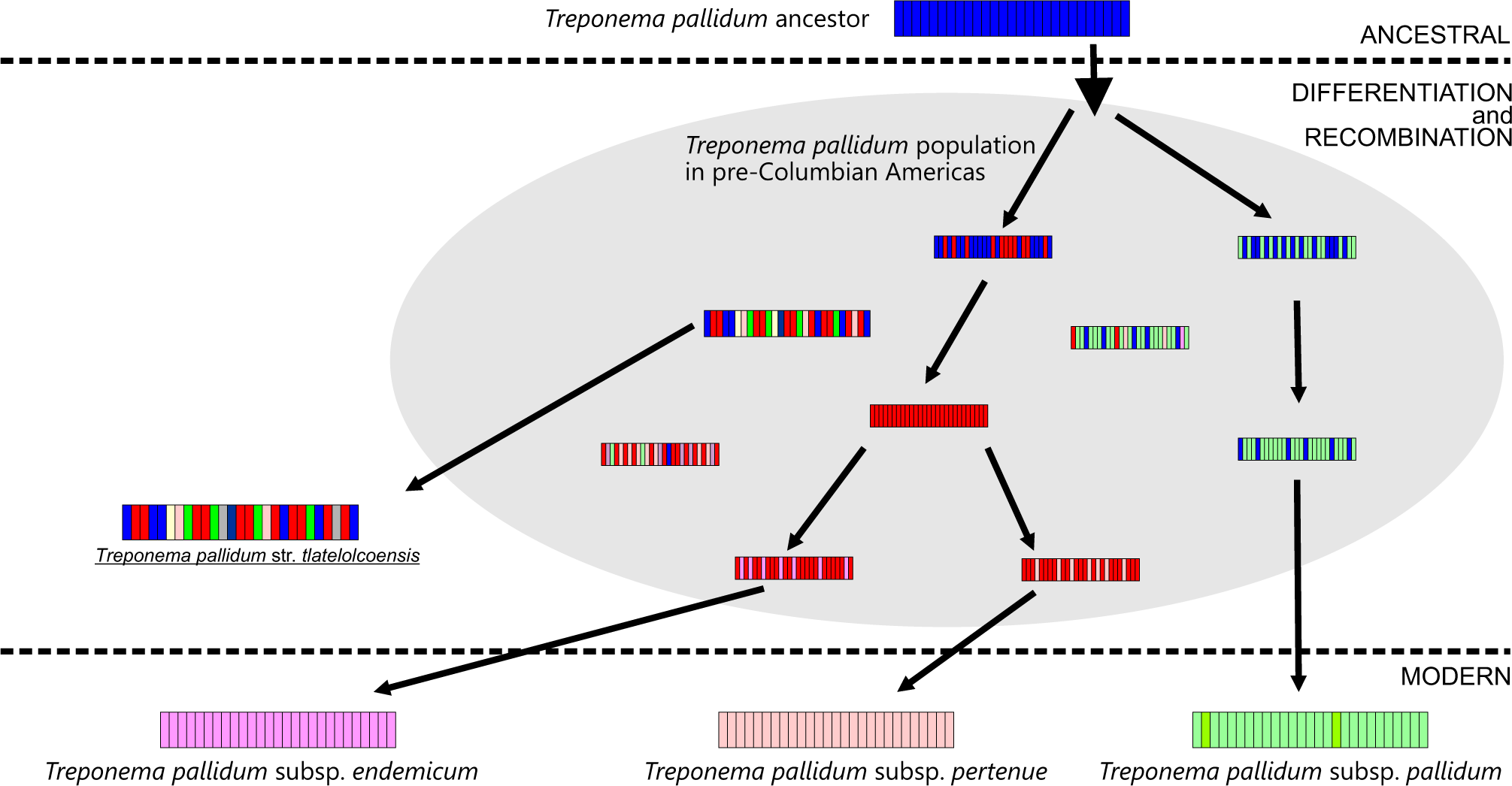
*T. pallidum* str. *tlatelolcoensis* belongs to a *Treponema* population that is ancestral to the three modern *T. pallidum* subspecies. The model of evolution has three stages (ancestral, differentiation and recombination, and modern) and uses the same colour scheme and segments as those of Fig. 2. In the upper part of the model is the ancestral *T. pallidum* genome. The central part of the model shows the emergence of the three *T. pallidum* subspecies through differentiation and recombination, as well as the presence of *T. pallidum* str. *tlatelolcoensis* in the population, together with two putative variants. The colour scheme uses the modern species as references and may give the impression that modern sequences did not recombine while some, like *T. pallidum* str. *tlatelolcoensis* did: this is arbitrary and would require more sequences from the ancestral population to reconstruct how the three modern subspecies were formed.

## Discussion

Mexico-Tlatelolco was a pre-Columbian city state built around 1337 on a small island of now dried-up Texcoco’s lake. It was regarded as the rival of the ancient capital Mexico-Tenochtitlan after Tlatelolco city developed a merchant empire considered one of the most important centers of activity of Mesoamerica^16^. Human sacrifice in pre-Columbian civilizations was highly ritualized and sacrificial inclusion was recently linked to impoverishment and to a high prevalence of infectious diseases^17^. It is thus noteworthy that the two children who carried the *T. pallidum* str. *tlatelolcoensis* characterized here were sacrificed girls, aged 4-6 years^15^, supporting the possibility that they displayed visible signs of the *Treponema* infection.

In the absence of previously reported paleomicrobiological traces of treponematoses in the New world^14^, the cases reported here represent the first genetic evidence of a *T. pallidum* complex species in pre-Columbian America. This result is in agreement with phylogenetic predictions^18^ and the observation of Pre-Columbian skeletal lesions characteristic of syphilis^19^. Importantly, this result is unambiguous as every single sequence read used for the genome assemblies belonged to the *T. pallidum* complex and the phylogenetic position of the resulting reconstructed genomes within the *T. pallidum* complex is confirmed by simple (NJ, parsimony) and more elaborated (ML) methods, which underlines the clear phylogenetic signal. Finally, while the two reconstructed genomes are incomplete, all analyses were restricted to the segments common to all the genomes used so as to avoid any potential bias due to sequence size differences. Taken together, these data thus clearly demonstrate the presence of a *T. pallidum* complex species in pre-Columbian America.

The pre-Columbian *T. pallidum* complex species identified in this study are closely related in global phylogenies to the modern *T. pallidum* complex species. In and of itself, this phylogenetic closeness is strong evidence that the two are direct descendants rather than just cousins. Indeed, any evolutionary scenario that would not make them direct descendants would require them to have separated well before the peopling of America, over 10kya^20^. This ancient split would be inconsistent with recent molecular estimates of the split between the three *T. pallidum* subspecies estimated at ∼4.6kya (with 95% highest posterior density intervals of ∼6.9-2.6 kya)^11^. Consistent with this, *T. pallidum* str. *tlatelolcoensis* includes both segments that are equivalent to those of two or three of the *T. pallidum* subspecies and segments that are equivalent to those of just one subspecies. This shows that pre-Columbian America is the region of the world where the transition from a single common ancestor to the direct ancestors of the three subspecies occurred. Hence the building blocks that are necessary to form the three *T. pallidum* subspecies were present in this geographical location and time period. 15^th^ century America is thus a required stop on the evolutionary path that led to modern *T. pallidum* subspecies. Notably, the largest group of segments related to just one *Treponema* subspecies in the genome of *T. pallidum* str. *tlatelolcoensis* is for *Treponema pallidum* subsp. *pallidum*, the subspecies that causes syphilis. And so, for the syphilis outbreaks to occur in Europe, those ∼75kb blocks (Supplementary Fig. 9a) would have had to be brought to Europe. Hence the characterization of *T. pallidum* str. *tlatelolcoensis* provides paleomicrobiological support to the hypothesis that Christopher Columbus ‘crew returning in Europe in 1493 brought venereal treponematosis to the continent, leading to the 1495 outbreak during the siege of Naples by the army of French King Charles VIII^2, 21^.

Interestingly, about half of the *T. pallidum* str. *tlatelolcoensis* genome has segments that are not related to just one single *T. pallidum* subspecies. While the exact age of the remains is not known and was initially associated with the Pre-Columbian era for this region of the Americas (1325-1520 CE), recent radiocarbon dating for remains excavated in the same locations points at date estimates that fall between 1332 and 1445 CE^17^. The strain characterized here may thus have preceded the arrival of Cristopher Columbus by 50-150 years. Hence the level of differentiation of *T. pallidum* strains in 1492 was more advanced than that observed for *T. pallidum* str. *tlatelolcoensis*, and full differentiation between the three *T. pallidum* subspecies likely occurred around the time of Cristopher Columbus’ travels. This raises interesting questions about how this differentiation is linked to the travels themselves (potential founder effect) or to the impact of encountering a naïve population in Europe.

## Methods

### Screening ancient human genomes for T. pallidum DNA

To identify studies that characterized ancient human genomes, we conducted bibliographical searches in Pubmed (https://pubmed.ncbi.nlm.nih.gov/) with ‘ancient’, ‘human’, ‘DNA’, and ‘genomes’ keywords. Raw genome data for the identified studies was obtained from the European Nucleotide Archive database (https://www.ebi.ac.uk/ena/browser/home). A total of 1,783 genomes from four regions of the world were obtained (Supplementary Figs. 1-2).

These ancient genomes were screened *in silico* using a low stringency approach with the Bowtie2 software^22^ and five probes: one specific to the *Treponema* genus (*flg*E) and four *T. pallidum*-specific ones (*pol*A, *tpp*47, *tpr*L and *tp*0619)^8, 9^. Specificity of the isolated reads was assessed through BLAST searches^23^ against the National Center for Biotechnology Information (NCBI) non-redundant nucleotide database.

### Reconstruction of the T. pallidum str. tlatelolcoensis genomes

After two positive cases were identified with our screening probes, we isolated *Treponema*-specific genomewide reads using a 3-step approach (Supplementary Fig. 3). To maximize the likelihood to capture all relevant reads, the first step was the same low stringency approach as that used for the screen with the probes using Bowtie2 software and a complete *T. pallidum* reference genome (*T. pallidum* subsp. *pallidum* strain Nichols; NC_021490.2).

A filtering stage was then performed, using Kraken2 software^24^ to assess specificity of the isolated reads and identify the source organisms for the *non-Treponema* reads. Monitoring of the results was performed with Krona software^25^. References for the five most represented *non-Treponema* genomes were obtained and a specificity analysis was conducted with Bowtie2 to isolate reads that were more related to the *Treponema* reference than to the *non-Treponema* references. Specificity was then reassessed with Kraken and the new five most represented *non-Treponema* genomes were used as negative references for another filtering. This loop was repeated until 25 negative genome references were used and the pool of reads reached 98% specificity for *Treponema*. In all these analyses a tolerance of eight differences comparing to the references was allowed.

Finally, to validate the specificity of the reads obtained after step #2, we conducted BLAST searches^23^ for each read as a third and final step. Searches were performed using the MEGABLAST program against the bacteria section of the NCBI non-redundant nucleotide database. Reads that produced no hit at this step were analyzed again using the BLASTN program. Reads with a non-*Treponema* best hit were discarded before the assembly step.

### Authenticity of ancient DNA

Authenticity of ancient DNA was verified by investigating for signs of cytosine deamination with mapDamage^26^ and for signs of DNA fragmentation by assessing read size distribution (Supplementary Fig. 4).

### Assembly and consensus

After filtering, the isolated reads were mapped against a *T. pallidum s*ubsp. *pallidum* strain Nichols genome sequence (NC_021490.2) using Mira assembly software^27^. Assemblies were visualized with the gap4 software of the STADEN package^28^ and a consensus sequence was extracted using a ‘Base frequencies’ algorithm with a 51% cutoff.

Representation of the coverage for the genome assembly was performed with Circleator^29^; in those representations, GC content and gene content is from the reference (*T. pallidum* subsp. *pallidum* strain Nichols).

### Phylogenetic analyses

*Treponema pallidum* str. *tlatelolcoensis* genomes were aligned with a representative set of 30 complete and modern *Treponema* genome sequences representing the three *T. pallidum* subspecies. Alignment was performed with the MAFFT software^30^ followed by manual corrections. In all analyses, the columns with alignment gaps or missing information were discarded (complete deletion datasets). Because of this, we did not include other ancient sequences: indeed, when trying to compare our sequences to those of Majander and colleagues^11^ for example, there was less than 10kb of common sequence.

All phylogenetic analyses were conducted with three methods: maximum-likelihood (ML), neighbor-joining (NJ) and parsimony. While ML methods are often the preferred choice over the other two methods, such a combination of simple and more elaborated phylogenetic methods can be helpful to detect underlying issues in the sequence data such as recombination, positions of functional divergence between paralogues, or biases created by outgroups as those issues often lead to incongruence in the results between the methods^31^.

NJ phylogenetic analyses were performed with MEGA11^32^ using the Tamura-Nei method with 500 replicates. PAUP*4.0a169^33^ and the tree bisection-reconnection branch swapping algorithm were used for parsimony analyses with 500 replicates and a heuristic search. ML analyses were performed with RAXML8^34^ under the available model (GTR+gamma) with 500 replicates (rapid bootstrapping).

Tree topology comparisons were performed using the Shimodaira-Hasegawa test of alternative phylogenetic hypotheses with re-sampling estimated log-likelihood optimization, and 10,000 bootstrap replicates (as implemented in PAUP*4.0b10). This comparison was made with the maximum likelihood model of DNA substitution defined using MODELTEST^35^ and the Akaike information criterion.

## Acknowledgements

M.G. benefited a Master grant from Fondation Méditerranée Infection, Marseille, France.

## Author contributions

M.D initiated the study. M.D and L.A-R designed and supervised the study. M.G conducted the study. M.G. and L.A-R performed the analyses. All authors worked on the data analysis and interpretation and wrote the manuscript.

## Competing interests

The authors declare no competing interest.

## Notes

### Competing Interest Statement

The authors have declared no competing interest.

## References

1. Crosby AW. The Early History of Syphilis: A Reappraisal. Am Anthropol 71, 218–227 (1969).

2. Tognotti E. The rise and fall of syphilis in Renaissance Europe. J Med Humanit 30, 99–113 (2009).

3. Arora N, et al. Origin of modern syphilis and emergence of a pandemic Treponema pallidum cluster. Nat Microbiol 2, 16245 (2016).

4. Rowley J, et al. Chlamydia, gonorrhoea, trichomoniasis and syphilis: global prevalence and incidence estimates, 2016. Bull World Health Organ 97, 548–562P (2019).

5. Fraser CM, et al. Complete genome sequence of Treponema pallidum, the syphilis spirochete. Science 281, 375–388 (1998).

6. Engelkens HJ, Vuzevski VD, ten Kate FJ, van der Heul P, van der Sluis JJ, Stolz E. Ultrastructural aspects of infection with Treponema pallidum subspecies pertenue (Pariaman strain). Genitourin Med 67, 403–407 (1991).

7. Giacani L, Lukehart SA. The endemic treponematoses. Clin Microbiol Rev 27, 89–115 (2014).

8. Gayet-Ageron A, Combescure C, Lautenschlager S, Ninet B, Perneger TV. Comparison of Diagnostic Accuracy of PCR Targeting the 47-Kilodalton Protein Membrane Gene of Treponema pallidum and PCR Targeting the DNA Polymerase I Gene: Systematic Review and Meta-analysis. J Clin Microbiol 53, 3522–3529 (2015).

9. Knauf S, et al. Gene target selection for loop-mediated isothermal amplification for rapid discrimination of Treponema pallidum subspecies. PLoS Negl Trop Dis 12, e0006396 (2018).

10. Giffin K, et al. A treponemal genome from an historic plague victim supports a recent emergence of yaws and its presence in 15(th) century Europe. Sci Rep 10, 9499 (2020).

11. Majander K, et al. Ancient Bacterial Genomes Reveal a High Diversity of Treponema pallidum Strains in Early Modern Europe. Curr Biol 30, 3788–3803 e3710 (2020).

12. Schuenemann VJ, et al. Historic Treponema pallidum genomes from Colonial Mexico retrieved from archaeological remains. PLoS Negl Trop Dis 12, e0006447 (2018).

13. Beale MA, Lukehart SA. Archaeogenetics: What Can Ancient Genomes Tell Us about the Origin of Syphilis? Curr Biol 30, R1092–R1095 (2020).

14. Harper KN, Zuckerman MK, Harper ML, Kingston JD, Armelagos GJ. The origin and antiquity of syphilis revisited: an appraisal of Old World pre-Columbian evidence for treponemal infection. Am J Phys Anthropol 146 Suppl 53, 99–133 (2011).

15. Morales-Arce AY, McCafferty G, Hand J, Schmill N, McGrath K, Speller C. Ancient mitochondrial DNA and population dynamics in postclassic Central Mexico: Tlatelolco (ad 1325–1520) and Cholula (ad 900–1350). Archaeol Anthropol Sci 11, 3459–3475 (2019).

16. The Oxford encyclopedia of Mesoamerican cultures. Oxford university press, Oxford New-York (2001).

17. Blevins KE, McGrane M, Mansilla Lory J, Guilliem Arroyo S, Buikstra JE. Structural Violence and Physical Death at Tlatelolco: Selecting the Chronically Malnourished for Sacrifice at a Late Postclassic Mesoamerican City (1300–1521 CE). Bioarchaeol int 7, 1–31 (2022).

18. Harper KN, et al. On the origin of the treponematoses: a phylogenetic approach. PLoS Negl Trop Dis 2, e148 (2008).

19. Tampa M, Sarbu I, Matei C, Benea V, Georgescu SR. Brief history of syphilis. J Med Life 7, 4–10 (2014).

20. Willerslev E, Meltzer DJ. Peopling of the Americas as inferred from ancient genomics. Nature 594, 356–364 (2021).

21. Baker BJ, Armelagos GJ. The origin and antiquity of syphilis: paleopathological diagnosis and interpretation. Curr Anthropol 29, 703–738 (1988).

22. Langmead B, Salzberg SL. Fast gapped-read alignment with Bowtie 2. Nat Methods 9, 357–359 (2012).

23. Altschul SF, et al. Gapped BLAST and PSI-BLAST: a new generation of protein database search programs. Nucleic Acids Res 25, 3389–3402 (1997).

24. Wood DE, Salzberg SL. Kraken: ultrafast metagenomic sequence classification using exact alignments. Genome Biol 15, R46 (2014).

25. Ondov BD, Bergman NH, Phillippy AM. Interactive metagenomic visualization in a Web browser. BMC Bioinformatics 12, 385 (2011).

26. Jonsson H, Ginolhac A, Schubert M, Johnson PL, Orlando L. mapDamage2.0: fast approximate Bayesian estimates of ancient DNA damage parameters. Bioinformatics 29, 1682–1684 (2013).

27. Chevreux B, et al. Using the miraEST assembler for reliable and automated mRNA transcript assembly and SNP detection in sequenced ESTs. Genome Res 14, 1147–1159 (2004).

28. Staden R, Beal KF, Bonfield JK. The Staden package, 1998. Methods Mol Biol 132, 115–130 (2000).

29. Crabtree J, Agrawal S, Mahurkar A, Myers GS, Rasko DA, White O. Circleator: flexible circular visualization of genome-associated data with BioPerl and SVG. Bioinformatics 30, 3125–3127 (2014).

30. Katoh K, Standley DM. MAFFT multiple sequence alignment software version 7: improvements in performance and usability. Mol Biol Evol 30, 772–780 (2013).

31. Abi-Rached L, Gilles A, Shiina T, Pontarotti P, Inoko H. Evidence of en bloc duplication in vertebrate genomes. Nat Genet 31, 100–105 (2002).

32. Tamura K, Stecher G, Kumar S. MEGA11: Molecular Evolutionary Genetics Analysis Version 11. Mol Biol Evol 38, 3022–3027 (2021).

33. Swofford DL. PAUP*: Phylogenetic analysis using parsimony (*and other methods), version 4.0. (eds. Sinauer, Sunderland, Massachusetts (2001).

34. Stamatakis A. RAxML version 8: a tool for phylogenetic analysis and post-analysis of large phylogenies. Bioinformatics 30, 1312–1313 (2014).

35. Posada D, Crandall KA. MODELTEST: testing the model of DNA substitution. Bioinformatics 14, 817–818 (1998).

