## Supplementary figures for "Pre-Columbian treponemes clarify worldwide spread of treponematosis"

| Region | Study | | Samples | | | Accession |
|---|---|---|---|---|---|---|
| | Authors | Year | # | Approximate age (years old) | Localisation | |
| Africa | Fregel et al | 2018 | 13 | 6,300 - 4,900 | Morocco, Iberia | PRJEB22699 |
| | Gallego Llorente et al | 2015 | 1 | 4,500 | Ethiopia | PRJNA295861 |
| | Lipson et al | 2020 | 2 | 8,000 - 3,000 | Cameroon | PRJEB32086 |
| | Rodriguez-Varela et al | 2017 | 5 | 1,400 - 900 | Canary Islands | PRJEB86458 |
| | Schlebusch et al | 2017 | 7 | 2,000 - 300 | South Africa | PRJEB22660 |
| | Schuenemann et al | 2017 | 3 | 3,400 -1,600 | Egypt | PRJEB15464 |
| | Skoglund et al | 2017 | 3 | 2,300 - 1,300 | South Africa | PRJEB21878 |
| Europe | Amorim et al | 2018 | 63 | 1,500 - 1,400 | Hungary and Italy | PRJNA433631 |
| | Brace et al | 2019 | 35 | 8,7500 - 2,500 | United Kingdom | PRJEB31249 |
| | Damgaard et al | 2018 | 30 | 3,200 - 600 | Europe | PRJEB20658 |
| | Furtwangler et al | 2020 | 96 | 6,500 - 3,700 | Switzerland, Germany, France | PRJNA608699 |
| | Gamba et al | 2014 | 13 | 7,800 - 2,800 | Hungary | PRJNA240906 |
| | Gelabert et al | 2021 | 1 | 25,000 | Georgia | PRJEB41420 |
| | Jarve et al | 2019 | 19 | 4,800 - 1,400 | Eastern Europe | PRJEB32764 |
| | Jensen et al | 2019 | 1 | 5,700 | Denmark | PRJEB30280 |
| | Kashuba et al | 2019 | 3 | 9,880 - 9540 | Sweden | PRJEB30667 |
| | Keller et al | 2012 | 1 | 5,300 | Italy | PRJEB2830 |
| | Krzewinska et al | 2018 | 23 | 1,100 - 900 | Sweden | PRJEB27220 |
| | Krzewinska et al | 2018 | 35 | 3,900 - 1,600 | Europe | PRJEB27628 |
| | Lamnidis et al | 2018 | 11 | 3,500 - 200 | NW Russia, Finland | PRJEB29360 |
| | Marcus et al | 2020 | 70 | 6,100 - 500 | Sardinia | PRJEB35094 |
| | Margaryan et al | 2020 | 442 | 4, 400 - 300 | Europe, Greenland | PRJEB37976 |
| | Mittnik et al | 2019 | 104 | 4,900 - 3,200 | Germany | PRJEB34400 |
| | Mittnik et al | 2018 | 38 | 9,500 - 2,200 | Northern Europe | PRJNA421333 |
| | Novak et al | 2021 | 38 | 6,200 | Croatia | PRJEB42243 |
| | Olalde et al | 2019 | 271 | 11,700 - 400 | Spain | PRJEB30874 |
| | Saag et al | 2017 | 10 | 6,300 - 4,500 | Estonia | PRJEB21037 |
| | Sanchez-Quinto et al | 2019 | 27 | 6,900 - 4,600 | Europe | PRJEB31045 |
| | Shroeder et al | 2019 | 15 | 5,000 | Poland | PRJEB28451 |
| | Skoglund et al | 2014 | 11 | 7,500 - 4,150 | Sweden | PRJEB6090 |
| | Veeramah et al | 2018 | 11 | 1,600 - 1,500 | Germany | PRJEB2307 |
| | Ziesemer et al | 2019 | 4 | 4,900 - 100 | Spain, Netherlands | PRJEB25624 |
| Asia | Damgaard et al | 2018 | 107 | 4,600 - 100 | Asia | PRJEB20658 |
| | Damgaard et al | 2018 | 74 | 11,500 - 500 | Asia | ERP107300, PRJEB26349 |
| | Feldman et al | 2019 | 8 | 15,600 - 8,700 | Turkey, Israel, Jordan | PRJEB24794 |
| | Fu et al | 2014 | 1 | 45,000 | Siberia | PRJEB6622 |
| | Kilinc et al | 2016 | 9 | 6,200 - 8,300 | Turkey | PRJEB14675 |
| | Omrak et al | 2015 | 2 | 4,800 - 6,700 | Turkey | PRJEB12155 |
| | Yang et al | 2017 | 1 | 40,000 | China | PRJEB20217 |
| America | Morales-Arce et al | 2019 | 13 | 1,100 - 500 | Mexico | PRJNA423230 |
| | Nakatsuka et al | 2020 | 20 | 5,800 - 100 | South Patagonia, Argentina | PRJEB39010 |
| | Nakatsuka et al | 2020 | 63 | 9,000 - 500 | Andes | PRJEB37446 |
| | Posth et al | 2018 | 3 | 11,200 - 600 | Central America | PRJEB28961 |
| | | | 46 | | South America | |
| | Raghavan et al | 2015 | 2 | 6,200 - 100 | North America | PRJEB9733 |
| | | | 6 | | Central America | |
| | | | 15 | | South America | |
| | Rasmussen et al | 2010 | 1 | 4,170 - 3,600 | Greenland | PRJNA46213 |
| | Rasmussen et al | 2014 | 1 | 12,700 - 12,550 | North America | PRJNA229448 |
| | Rasmussen et al | 2015 | 1 | 8,340 - 9,200 | North America | PRJNA284124 |
| | Ziesemer et al | 2019 | 2 | 1,000 - 600 | Guadeloupe | PRJEB25624 |
| | | | 2 | | USA | |

**Supplementary Figure 1**. Ancient genomes investigated in the study. Genomes are organized by geographical region and the following information is given for each line: study information (authors and year), sample information (approximate age and localisation), and accession number for the sequences.

| Region | Genomes (#) | Proportion (%) |
| --- | --- | --- |
| Africa | 34 | 1.9 |
| Europe | 1,372 | 76.9 |
| Asia | 202 | 11.3 |
| America | 175 | 9.8 |
| Total | 1,783 | 100 |

**Supplementary Figure 2**. Total number of ancient genomes investigated in the study for each region of the world.

**a**

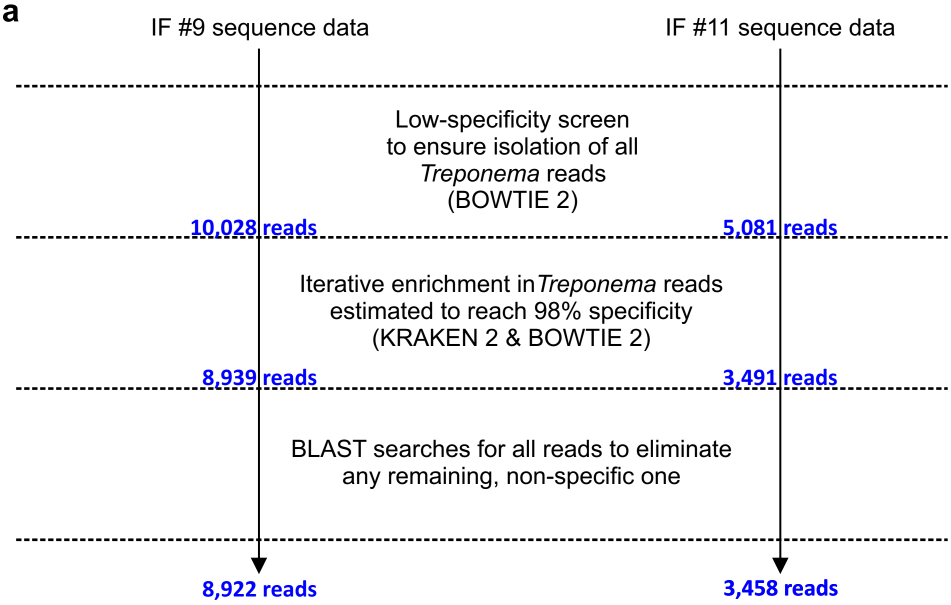


**b**

| | Reads # | | | | | | | |
|---|---|---|---|---|---|---|---|---|
| | From KRAKEN2 analysis | Top hit in BLAST searches | | | | | Summary | |
| | | *T. pallidum* | *T. paraluisleporis* *T. paraluiscuniculi* | *T. phagedenis* | Non *Treponema* | No significant similarity | *Trepanoma* specific | Non specific |
| *IF #9* | 8,939 | 8,860 | 60 | 2 | 16 | 1 | 8,922 | 17 |
| *IF #11* | 3,491 | 3,440 | 17 | 1 | 23 | 10 | 3,458 | 33 |

**Supplementary Figure 3**. Isolation of the *Treponema*-specific reads from the sequence data of the two positive individuals. (**a**) Three-step procedure used to isolate the *Treponema*-specific reads from the sequence data for individuals IF #9 and IF #11. For each step and individual, the number of reads isolated is given in blue. (**b**) Results of the filtering used in the third step. The reads isolated at the end of this step were used for the genome assemblies.

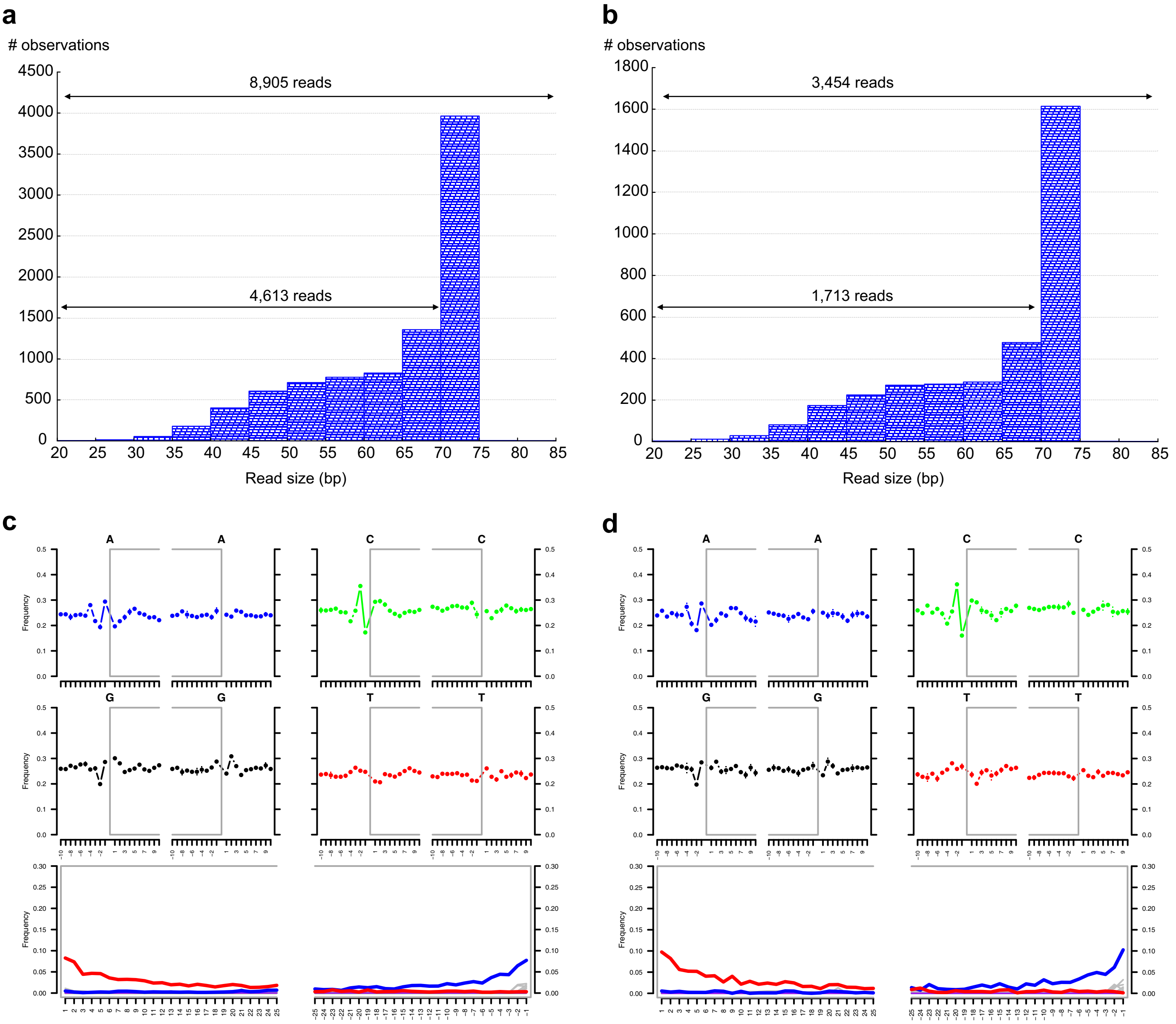


**Supplementary Figure 4**. Reads used for the *T. pallidum* str. *tlatelolcoensis* genome reconstructions have the characteristics of ancient DNA. (**a** and **b**) Size distribution for the 8,905 reads isolated from individual IF #9 and used in the genome assembly (**a**) and for the 3,454 reads isolated from individual IF #11 and used in the genome assembly (**b**). As is expected for ancient DNA, fragments are degraded and ~50% of the reads are <70bp: 4,613 reads out of 8,905 for IF #9 (52%) and 1,713 reads out of 3,454 for IF #11 (50%). Overrepresentation for the 70-75bp category is due to the 80bp sequencing that was performed and the clustering of all the fragments >70bp in just one category. (**c** and **d**) mapDamage analysis for the reads isolated from individual IF #9 (**c**) and for the reads isolated from individual IF #11 (**d**). The four upper mini-plots show base frequencies in and around the reads. The lower plot shows the position-specific substitutions. As is expected for ancient DNA, an excess of C to T substitutions is observed in the 5' end (left, red line) and of G to A substitution in the 3' end (right, blue line).

**a**

| | | Estimated remaining cytosine deamination (%) | *T. pallidum* str. *tlatelolcoensis* #1 vs #2 | | |
|---|---|---|---|---|---|
| | | | Differences (#) | Overlap (bp) | % divergence |
| Raw reads | | 100.0 | 1,104 | 76,901 | 1.44 |
| Map-damage corrected reads | cut 5 | 56.2 | 665 | 58,447 | 1.14 |
| | cut 10 | 35.0 | 438 | 40,419 | 1.08 |
| | cut 15 | 20.4 | 232 | 23,724 | 0.98 |

**b**

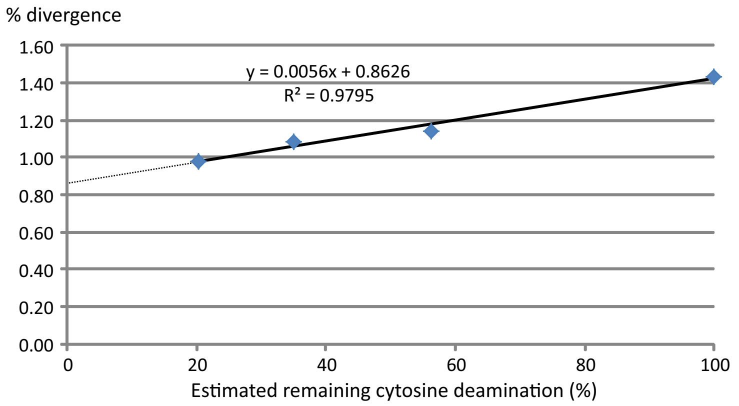


**Supplementary Figure 5**. Comparison of the two *Treponema pallidum* str. *tlatelolcoensis* genomes. (**a**) The two *Treponema pallidum* str. *tlatelolcoensis* genomes were each assembled either from raw reads or from reads whose ends were trimmed by 5bp ('cut 5'), 10bp ('cut 10'), or 15bp ('cut 15'), and the resulting consensus sequences were compared. For each comparison, the estimated remaining amount of cytosine deamination (as assessed by MapDamage) is given together with the number of differences, the size of the overlapping regions and the overall divergence (as a percentage) between the two genomes. (**b**) Correlation between the percentage of divergence between the two genomes and the estimated remaining level of cytosine deamination. The intersection between the trend line and the vertical axis occurs at 0.86, suggesting that when all the differences due to cytosine deamination are removed, the divergence between the two genomes will be 0.86%.

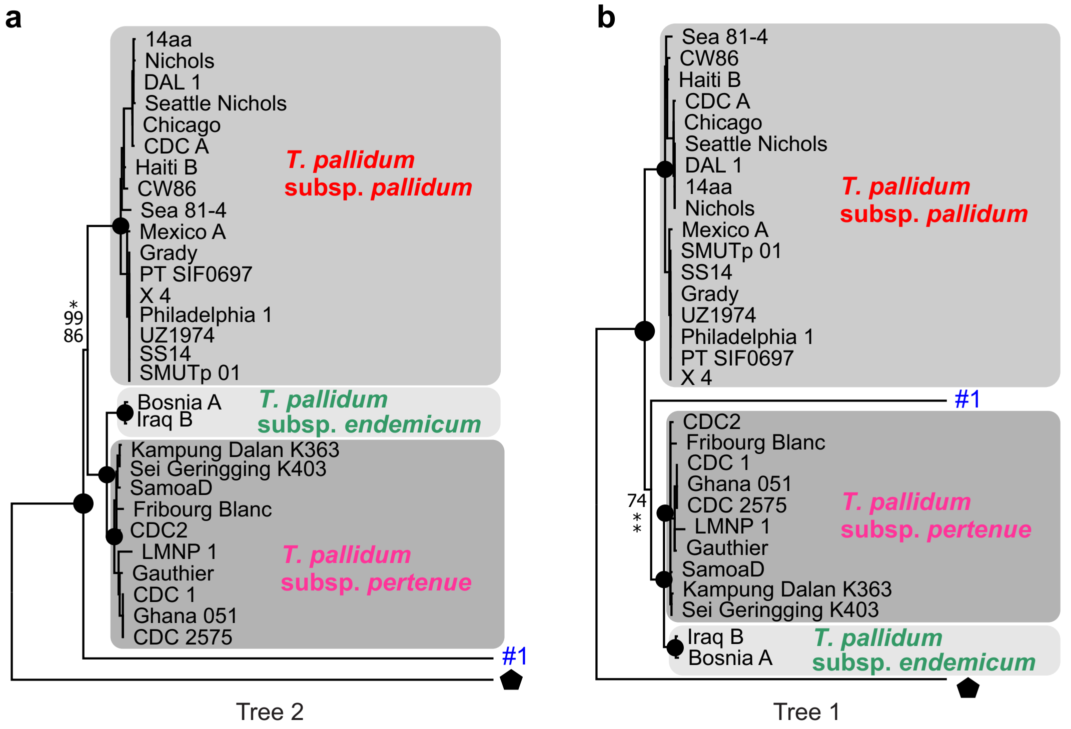


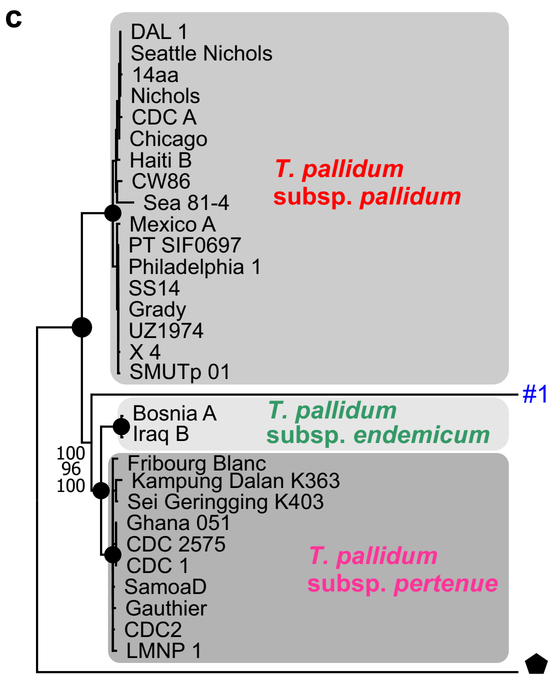


**d**

Shimodaira-Hasegawa test:
SH test using RELL bootstrap (one-tailed test)
Number of bootstrap replicates = 10000

| Tree | -ln L | Diff -ln L | P |
|---|---|---|---|
| 1 | 304700.94211 | (best) | |
| 2 | 304720.83761 | 19.89550 | 0.0113* |

* P < 0.05

**Supplementary Figure 6**. Significant incongruence between the phylogenetic methods. (**a** and **c**) The dataset used in Fig. 1f was partitioned in two halves and each half was analysed with three phylogenetic methods, as described in Fig. 1f. For the 'left' side of the dataset, differences were observed between the NJ and parsimony analyses (**a**) and the ML analysis (**b**). For the 'right' side of the dataset, all three methods yielded the same topology (**c**). (**d**) The Shimodaira-Hasegawa test of alternative phylogenetic hypotheses shows that the difference in topology between the phylogenetic trees of panels (**a**) and (**b**) is significant (alpha=0.05).

| *T. pallidum* subspecies | | Average number of differences | | | | | | | | Average segment required for 10 differences (kb) | | |
|---|---|---|---|---|---|---|---|---|---|---|---|---|
| # 1 | # 2 | # (+/- SE) | | 95% CI | | | per kb | | | | | |
| endemicum | pertenue | 792 | (22) | 749 | - | 835 | 0.66 | - | 0.74 | 15.1 | - | 13.5 |
| endemicum | pallidum | 1,738 | (34) | 1,671 | - | 1,805 | 1.48 | - | 1.60 | 6.8 | - | 6.3 |
| pertenue | pallidum | 1,662 | (34) | 1,595 | - | 1,729 | 1.41 | - | 1.53 | 7.1 | - | 6.5 |

**Supplementary Figure 7**. *T. pallidum* subspecies comparisons to establish the size of the windows to use in the phylogenetic scan. Comparisons between *T. pallidum* subspecies were performed to determine the genomewide average number of differences. From these data, we estimated the average size of the genomic segments required to have 10 differences or more between subspecies.

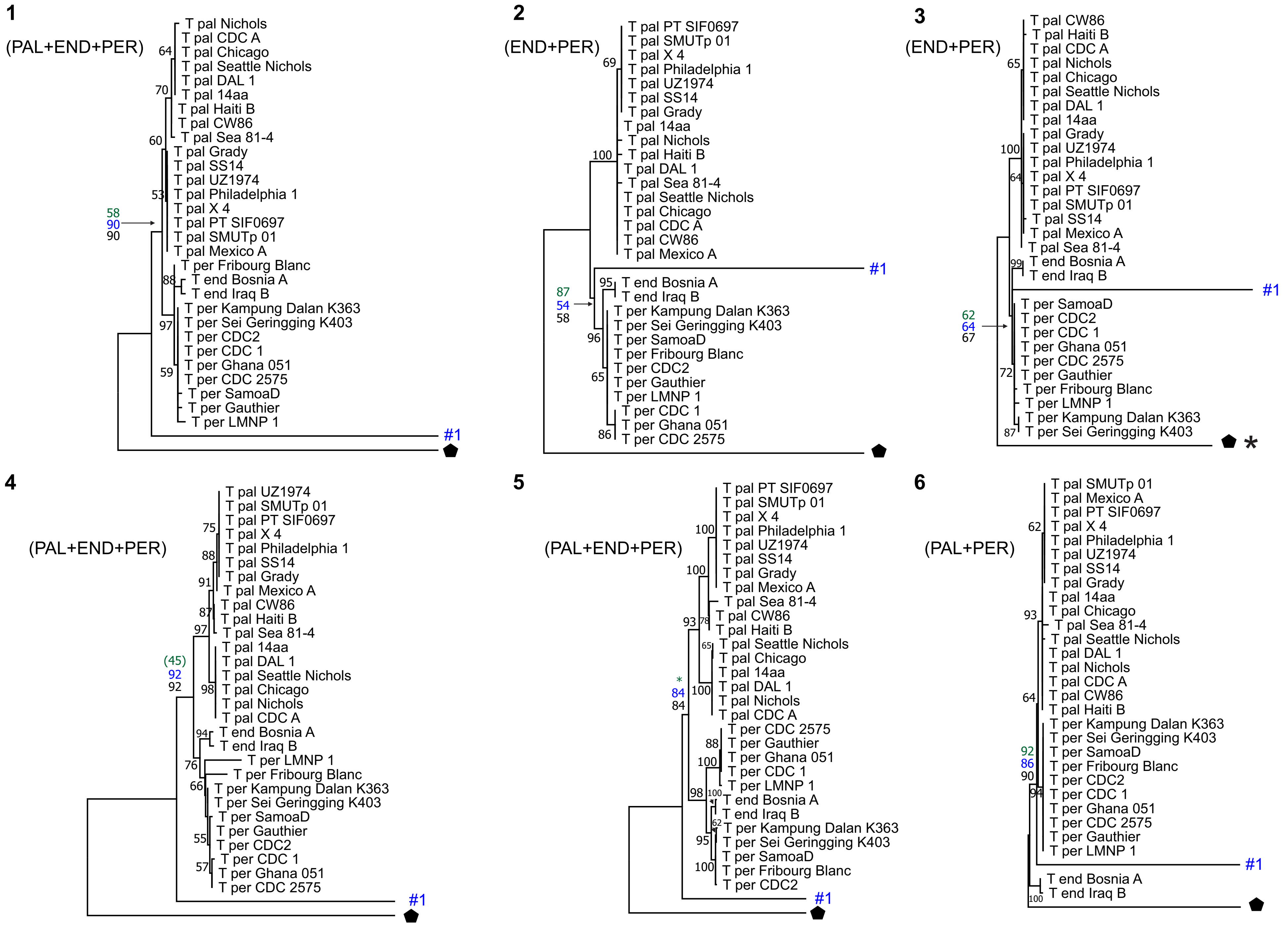


Supplementary Fig. 8

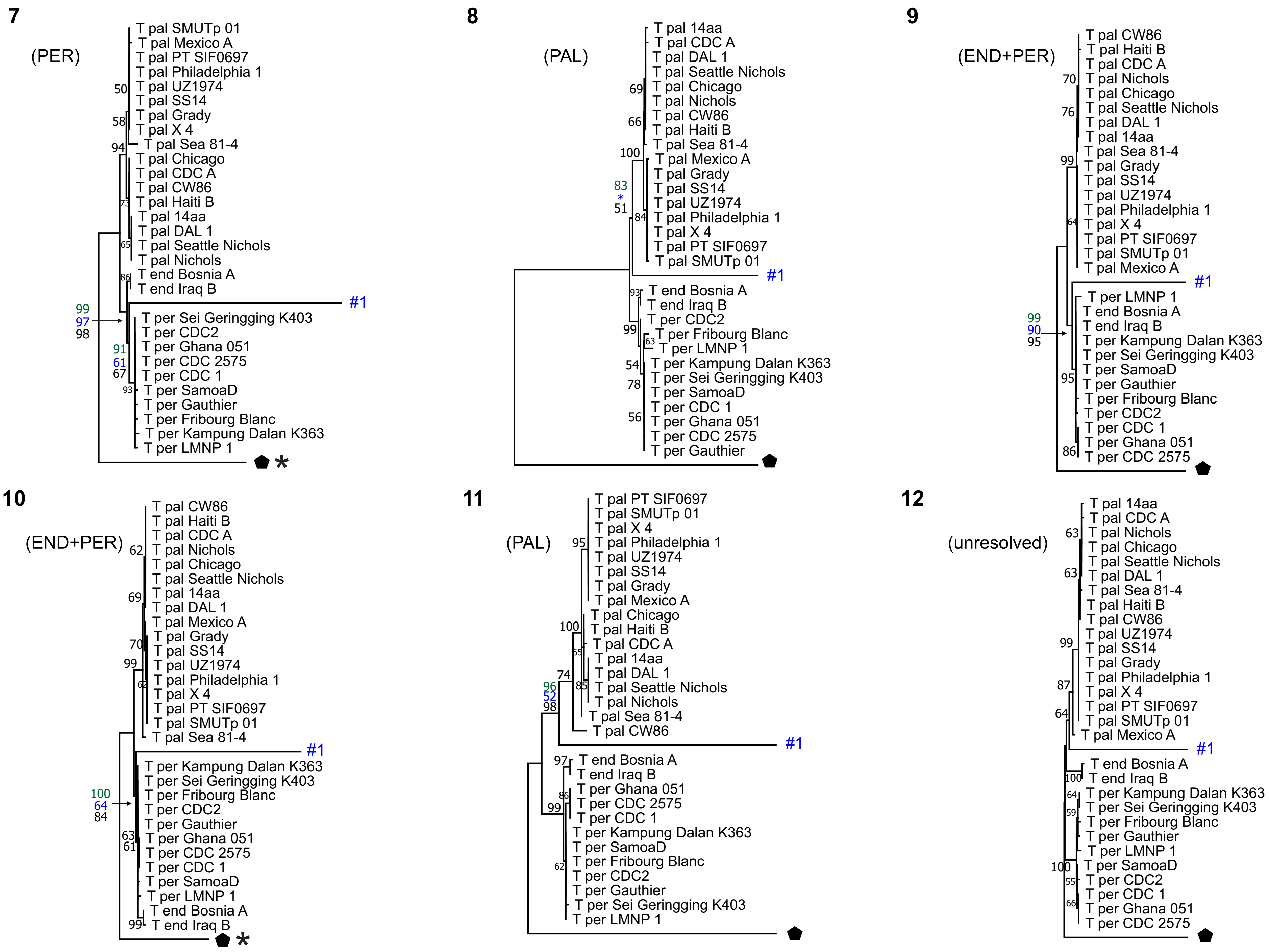


Supplementary Fig. 8

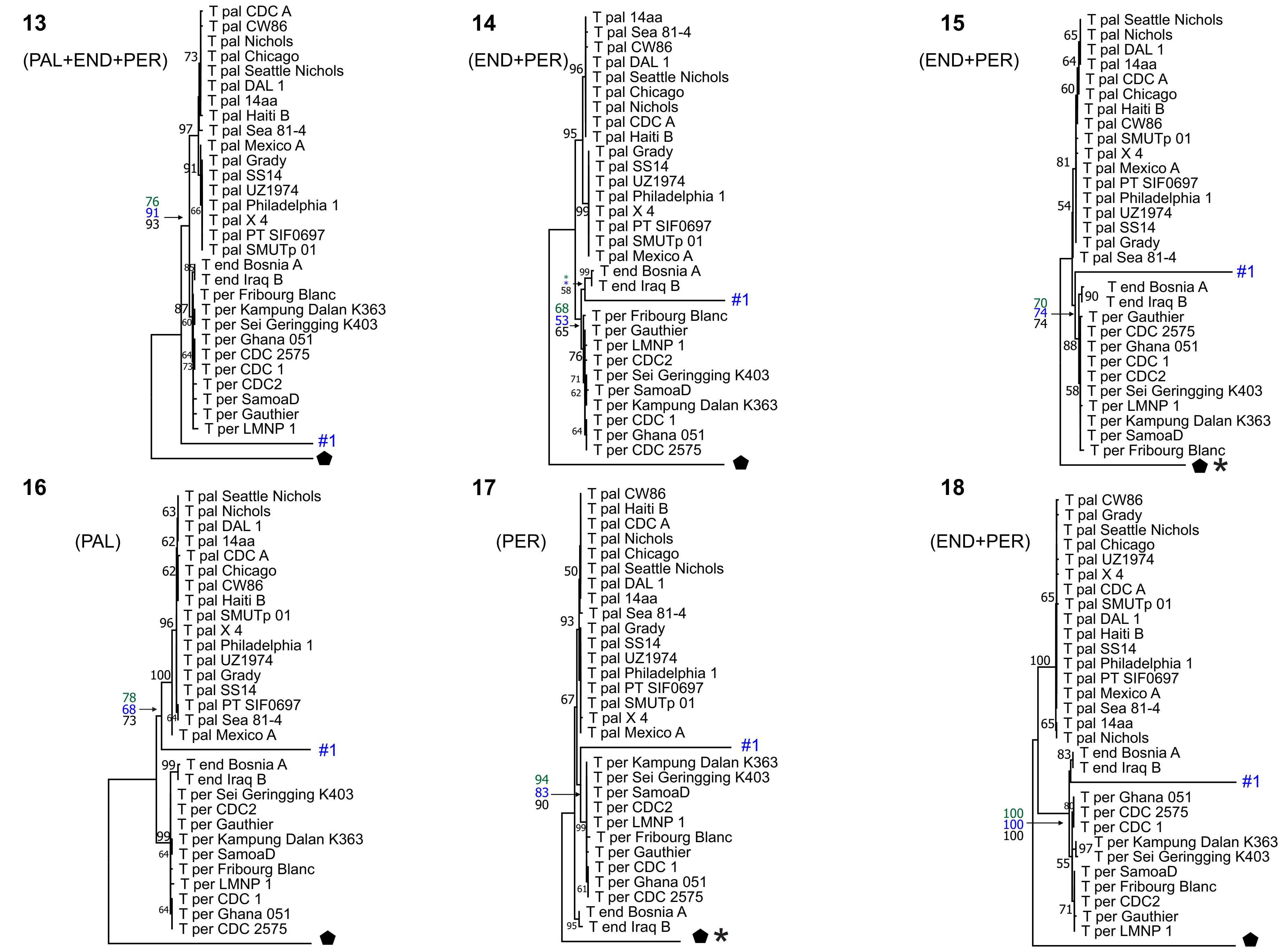


Supplementary Fig. 8

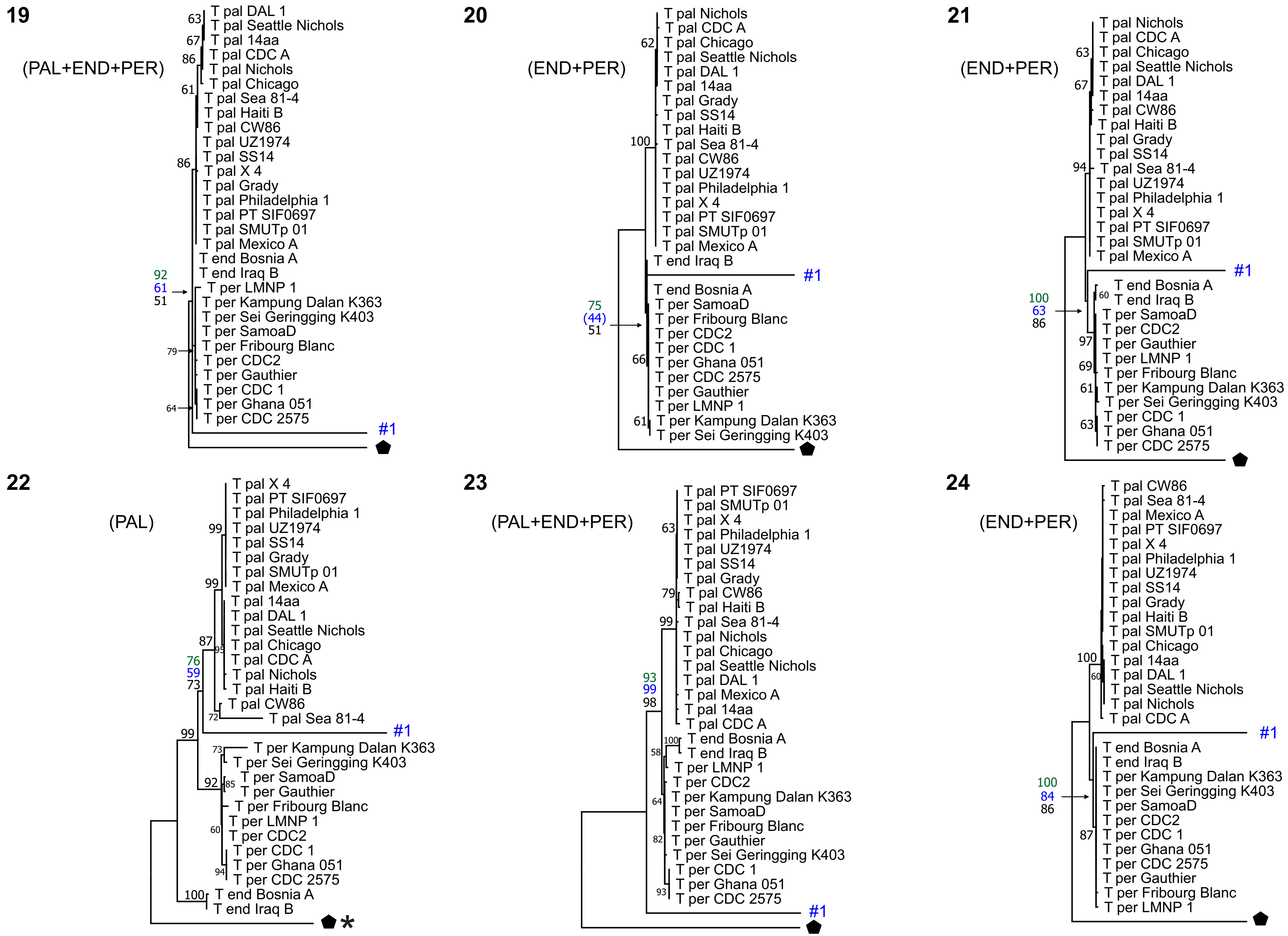


Supplementary Fig. 8

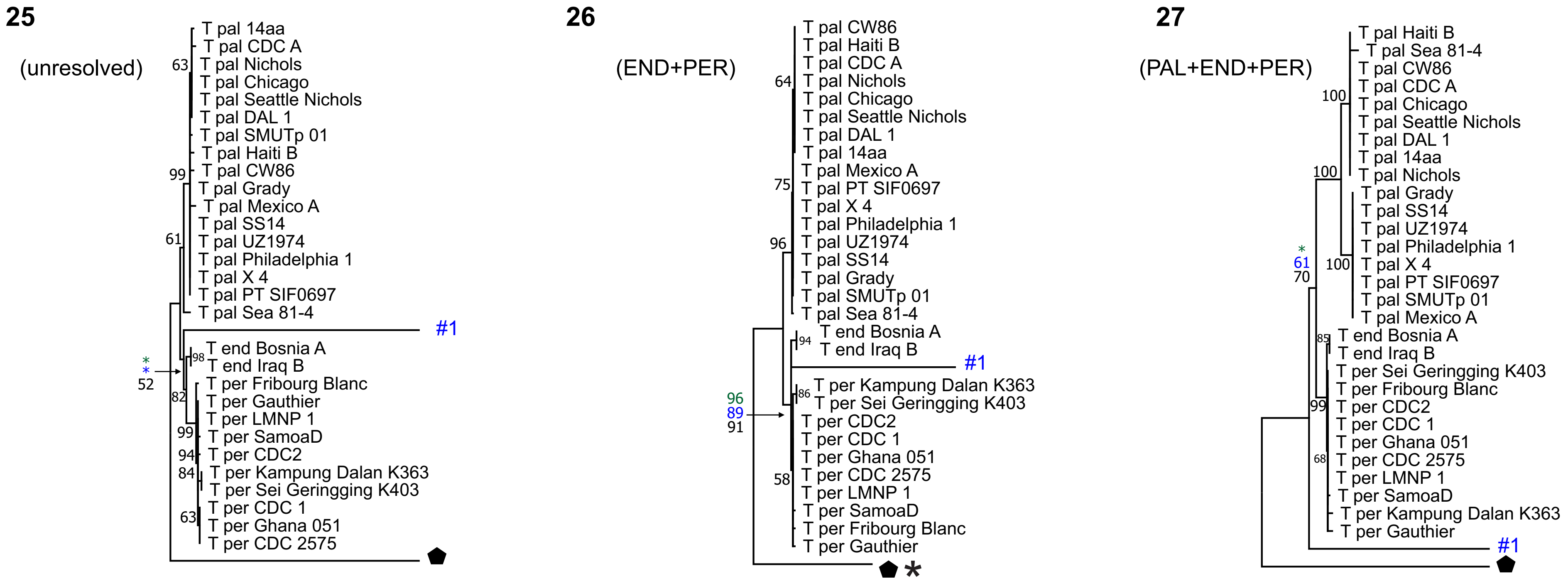


**Supplementary Figure 8**. Phylogenetic scan for the Tlatelolco genome. The *T. pallidum* str. *tlatelolcoensis* genome #1 was divided in 27 segments of 15kb each and phylogenetically compared to the corresponding sequences of 30 other *Treponema* genomes using NJ, parsimony, and ML methods. NJ tree topologies with midpoint rooting are used for the display. NJ bootstrap support is given when >50; ML and parsimony support is given only for the node relevant for the position of the *T. pallidum* str. *tlatelolcoensis* sequence in the tree (from top to bottom: NJ, parsimony, ML; bootstrap support <50 is given in parenthesis; *, clustering not supported). For each of the 27 segments, the relationship between *T. pallidum* str. *tlatelolcoensis* and the other three *T. pallidum* subspecies is summarized in parenthesis under the segment number using the following abbreviations: PAL (*T. pallidum* subsp. *pallidum*), END (*T. pallidum* subsp. *endemicum*), PER (*T. pallidum* subsp. *pertenue*). For example, '(PAL+END+PER)' means that *T. pallidum* str. *tlatelolcoensis* is clustering as a sister group to all three *T. pallidum* subspecies. Black pentagons, *T. paraluiscuniculi* outgroup (asterisks indicate that the root was forced with the outgroup).

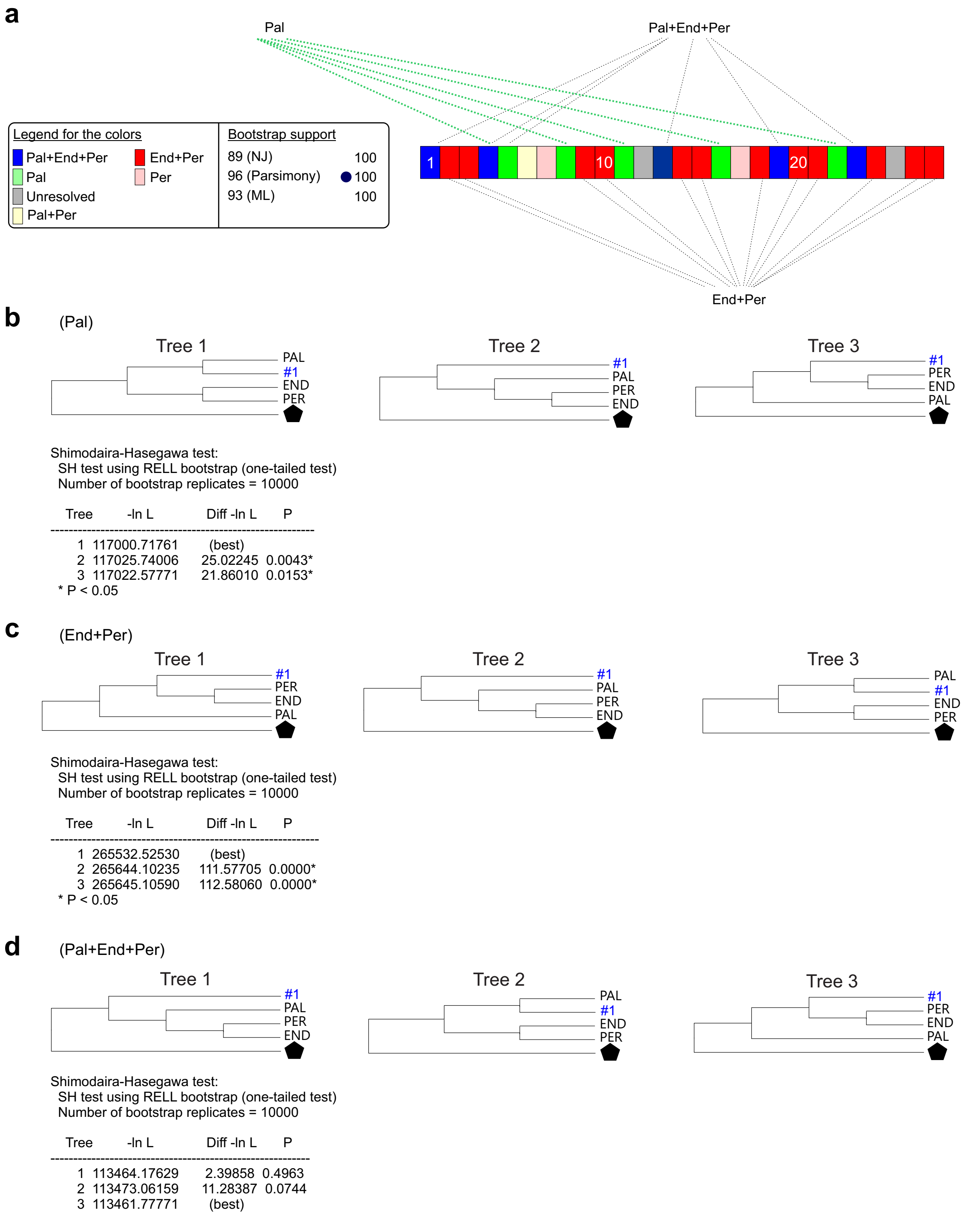


**Supplementary Figure 9**. Tree topology comparisons. (**a**) ML version of Fig.2 with differences for segments #5 and #27 (see Supplementary Fig. 8). (**b** and **d**) Tree topology comparisons for the 'Pal' (**b**), 'End+Per' (**c**), or 'Pal+End+Per' (**d**) segments displayed in (**a**). For each comparison, the three topologies compared are given, together with the results of the Shimodaira-Hasegawa test.
